# Structural basis for the inhibition of *Trypanosoma brucei* enolase by a camelid single-domain antibody

**DOI:** 10.64898/2025.12.17.694860

**Authors:** Zeng Li, Natalia Smiejkowska, Julie Vansevenant, Jimmy Mertens, Pieter Van Wielendaele, Joar Esteban Pinto Torres, Stefan Magez, Yann G.-J. Sterckx

## Abstract

*Trypanosoma brucei* is an extracellular protozoan that causes neglected tropical diseases in both humans and livestock. The parasite has a bipartite life cycle involving an insect vector and a mammalian host. Within the latter, it mainly thrives as a blood-borne parasite that relies on glycolysis to support its energy metabolism. It is for this reason that trypanosomal glycolytic enzymes have been investigated as potential targets for the development of trypanosome-killing drugs. Recent work from our lab has shown that they are also interesting biomarkers for the detection of active trypanosome infections. *T. brucei* enolase (*Tbr*ENO) is a trypanosomal glycolytic enzyme that has gathered interest in both drug and diagnostics development. In this paper, we report the discovery of a camelid single domain antibody (sdAb aka nanobody) that specifically recognises and inhibits *Tbr*ENO. The sdAb’s inhibitory mechanism is unraveled through a combination of protein biochemistry, biophysics, and structural biology.

**Author summary:** *Trypanosoma brucei* is a unicellular parasite that lives in the bloodstream of humans and animals, where it causes serious but often overlooked diseases. Because it depends heavily on breaking down glucose to produce energy, the parasite’s glucose-processing proteins (called glycolytic enzymes) have become important targets for both new treatments and improved diagnostic tools. One of these proteins, called *T. brucei* enolase (*Tbr*ENO), has recently drawn attention for its potential in drug development and disease detection. In this study, we discovered a special type of antibody, known as a camelid single-domain antibody (sdAb aka nanobody), that can specifically recognize and block the activity of *Tbr*ENO. We employed a combination of various laboratory techniques to understand exactly how the sdAb binds to and inhibits the enzyme. Our findings provide new insight into how *Tbr*ENO can be inhibited in a way that does not require active site binding and highlight the value of sdAbs as precise tools for targeting key parasite proteins.

## Introduction

Trypanosomiasis remains a major health and socio-economic concern across large areas of the (sub)tropics. The causative agents are protozoan parasites of the *Trypanosoma* genus. Depending on the species, African trypanosomes cause diseases in humans (*T. brucei gambiense* and *T. brucei rhodesiense*) or livestock (*T. congolense, T. vivax, T*.*brucei evansi, T. brucei equiperdum* and *T. brucei brucei* ) known as human and animal African trypanosomiasis (HAT and AAT, respectively) [1]. *T. brucei evansi* is the most widespread AAT-causing trypanosome as it is mechanically transmitted by a variety of hematophagous biting flies, which has enabled its spread beyond the African “tsetse belt” [2, 3]. The parasite infects both livestock and wildlife (e.g. horses, cattle, camels, and buffalos) in which it causes an AAT known as “surra” or “mal de caderas”, a severe pathology characterized by weight loss, drastic reductions of draft power, diminished meat and milk production that often results in the death of infected animals [4, 5].

Although rare, atypical HAT cases caused by *T. brucei evansi* have recently been reported, suggesting that the parasite may be an emerging zoonotic pathogen [6]. Compared to tsetse-transmitted AAT, controlling “surra” is more challenging since *T. brucei evansi* lacks vector specificity. Therefore, disease control is largely based on the use of trypanocidal drugs and the prevention of infection [3]. Drug treatment in livestock should ideally be combined with a conclusive diagnosis of infection or cure.

Within this context, we described the development of nucleic acid [7] and antigen-based [8] detection methods. For the latter, the glycolytic enzyme *T. brucei evansi* enolase (ENO) was identified as a potential diagnostic biomarker through an unbiased approach employing camelid single-domain antibodies (sdAbs aka nanobodies). Trypanosomal ENO displays a 100% amino acid sequence conservation within *T. brucei* subspecies (called *Tbr*ENO from hereon) and was previously investigated as a target for the development of novel trypanocidal drugs [9–12].

ENOs (aka phosphopyruvate hydratases) are some of the most abundantly expressed cytosolic proteins in many organisms and are highly conserved across all domains of life [13]. They primarily function as enzymes in glycolysis and gluconeogenesis through the reversible interconversion of D-2-phosphoglycerate to phosphoenolpyruvate (PEP). However, in addition to their metabolic function, ENOs also perform so-called “moonlighting” functions related to (patho)physiological processes in different organisms [14]. Interestingly, various pathogens appear to employ ENO at the cell surface as a receptor for host plasminogen (PLG), which is proposed to enable the crossing of cell barriers, thereby contributing to pathogen invasiveness and migration within the host [15–24]. The availability of high-resolution crystal structures of ENOs from various organisms has provided a detailed understanding of their structure-function relationship and catalytic mechanism [9–11, 25–41]. In most organisms, ENOs are typically homodimers, although the existence of functional heterodimers and homo-octamers has been reported [33, 42–44]. The structure of the ENO monomer consists of a small N-terminal *α* + *β* domain composed of four *α*-helices and three *β*-strands, followed by a C-terminal eightfold *α*/*β* barrel domain displaying a *ββαα*(*βα*)_6_ topology. The active site lies at the interface of both domains (Figure 1A) and accommodates both the substrate/product and two divalent cations (Mg^2+^ or Zn^2+^, with a preference for Mg^2+^) required for catalysis. The binding of these ligands to the active site and their subsequent release after catalysis is sequential. A first Mg^2+^ ion binds to ‘site I’ formed by a triad of highly conserved residues (Asp243, Glu291, and Asp318 for *Tbr*ENO). This is necessary for subsequent substrate binding to the active site: while the ‘site I’ Mg^2+^ accommodates the substrate’s carboxyl group, *Tbr*ENO residues (Ser40, His156, Gln164, Glu165, Glu208, Leu341, Lys343, Arg372, Ser373, Lys394) interact with the remainder of the molecule. Next, the bound substrate enables binding of a second Mg^2+^ ion to ‘site II’ formed by an active site residue (Ser41) and the substrate’s phosphoryl and carboxyl groups. After catalysis, product release requires the dissociation of the ‘site II’ Mg^2+^ ion, which explains why high concentrations of divalent metal ions have an inhibitory effect on enzyme activity [11]. Substrate binding, catalysis, and product release rely on the structural plasticity of three regions surrounding the active site, which are termed ‘loops’ despite containing secondary structure elements (Figure 1B): ‘loop 1’ (Ser37 to Tyr44), ‘loop 2’ (Val150 to Phe166), and ‘loop 3’ (Cys244 to Glu272). These loops adopt various conformations ranging from ‘open’ to ‘closed’ (and intermediates thereof, often called ‘semi-open/closed’). Mechanistically, ENOs employ a general acid-base catalysis for which there are two proposals. While in the first mechanism His156 (part of ‘loop 2’) and Lys343 act as the general acid-base pair [45], the second (more widely accepted) proposal features Glu208-Lys343 as the general acid-base pair [46]. In the latter, His156 plays a role in transition state stabilization through charge complementation. In any case, a correctly positioned His156 is required to obtain a functional active site and thus a catalytically competent enzyme. The highly conserved dimerization interface, monomer subunit structure, and especially the active site bestows ENOs from distinct species with similar kinetic properties.

**Fig 1.**
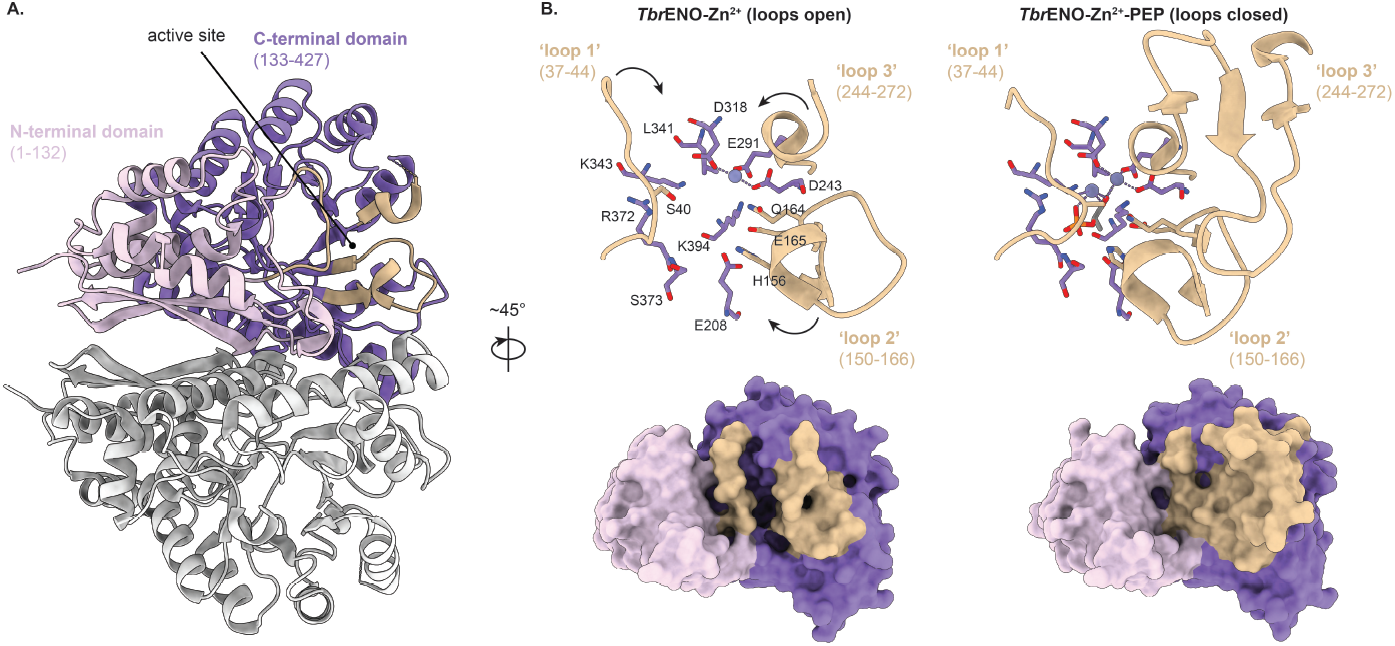
Structural basis for ENO enzymatic activity. (A.) Cartoon representation of the *Tbr*ENO dimer structure. While one *Tbr*ENO dimer monomer is colored light gray, the domains of the other monomer are color-coded and the domain boundaries are shown. The position of the enzyme’s active site at the domain interface is indicated for the reader’s convenience. (B.) Comparison of *Tbr*ENO structures with different active site loop conformations: *Tbr*ENO-Zn^2+^ and *Tbr*ENO-Zn^2+^-PEP (loops open and closed; PDB IDs 2PTW and 2PTY, respectively [11]). The top panels show a close-up of loops 1, 2, and 3 (cartoon representation) and the active site residues (shown in stick representation). The bottom panels display a surface representation of the *Tbr*ENO structures, which visualises active site accessibility depending on the loop conformation.

In this paper, we present the structural basis for the inhibition of *Tbr*ENO by a camelid sdAb (sdAbR1-10). Enzymatic activity assays reveal that sdAbR1-10, which was originally identified within a diagnostic context [8], is a potent *Tbr*ENO inhibitor. The *Tbr*ENO - sdAbR1-10 interaction was further characterized by analytical size exclusion chromatography (SEC), isothermal titration calorimetry (ITC) and macromolecular X-ray crystallography (MX). sdAbR1-10 recognizes an epitope containing *Tbr*ENO ‘loop 2’ and ‘loop 3’ residues, which prevents these loops from fully closing, thereby impairing *Tbr*ENO catalytic activity. Residue conservation analysis of the targeted epitope reveals why sdAbR1-10 efficiently recognizes trypanosomal ENOs, but not human and bovine ENOs.

## Materials and methods

### Recombinant protein production and purification

The generation of sdAbR1-10 by alpaca immunization, the recombinant production of C-terminally His-tagged sdAbR1-10 and N-terminally His-tagged *Tbr*ENO in *E. coli*, and subsequent purification by immobilized metal affinity chromatography (IMAC) and size exclusion chromatography (SEC) have previously been described [8].

### Analytical SEC

The stoichiometry of the *Tbr*ENO - sdAbR1-10 complex was performed by analytical SEC using an ENrich SEC 650 10 x 300 Column (Bio-Rad), pre-equilibrated in buffer A (50 mM sodium phosphate, 5 mM MgCl_2_, pH 7.0). Samples of 500 *µ*l containing 500 *µ*g recombinant *Tbr*ENO mixed with sdAbR1-10 at various molar ratios expressed in *Tbr*ENO monomer equivalents (*Tbr*ENO:sdAbR1-10 ratios of 2:1, 2:2, 2:3, and 2:4 respectively) were incubated for 1 h on ice prior to application onto the column, followed by sample elution at a flow rate of 1 ml min^-1^. The column was calibrated with the BioRad molecular mass standard under the same conditions. The elution peaks of all chromatograms were analyzed by SDS-PAGE.

### Isothermal titration calorimetry

The interaction between *Tbr*ENO and sdAbR1-10 was investigated by isothermal titration calorimetry (ITC) on a MicroCal PEAQ-ITC instrument (Malvern). sdAbR1-10 (100 *µ*M) was titrated into the sample cell containing *Tbr*ENO (10 *µ*M). Both proteins were extensively dialyzed against the same buffer (50 mM sodium phosphate, 5 mM MgCl_2_, pH 7.0) to exactly match the buffer composition. Before being examined in the calorimeter, all samples were degassed for 10 min at a temperature close to the titration temperature (25^°^C) to prevent long equilibration delays. The reference power was set to 5 *µ*cal s^-1^ and a stirring speed of 750 rpm was used. An equilibrium delay of 180 s before the start of each measurement was employed, while a spacing of 150 s between each injection was used. Twelve injections with constant volume (3.0 *µ*L) were performed during data collection. The first injection was always 0.4 *µ*L and its associated heat was never considered during data analysis. To determine the injection heats, control titrations were performed consisting of sdAbR1-10 injections into the buffer-filled cell (thus in the absence of *Tbr*ENO). Baseline adjustment, control subtraction, and data analysis were performed using the MicroCal PEAQ-ITC analysis software. The data were analyzed with the “one set of sites” binding model resulting in fitted values for the stoichiometry of the interaction (N), the equilibrium dissociation constant (K_*D*_), and the change in enthalpy (ΔH) and entropy (ΔS) associated with the binding events. All experiments were performed in triplicate.

### Inhibition assay

The enzymatic assay was performed as previously described [8], with the following modifications to evaluate the inhibitory properties of sdAbR1-10. Briefly, the inhibition assay was performed at 25^°^C in a 100 *µ*l reaction mixture containing 0.1 M triethanolamine/HCl, pH 7.6, 1.1 mM ADP (Sigma A2754), 0.42 mM NADH (Sigma N8129), 2 mM MgSO_4_, and 17 mM KCl. The auxiliary enzymes PYK and LDH mixture solution (Sigma P0294) were used at 4 and 6 U ml^-1^. The substrate 2-PGA (Sigma 79480) was added in varying concentrations between 0 and 1 mM as indicated.

The inhibitory properties were evaluated by adding sdAbR1-10 in variant concentrations between 0 to 2 mM.

### Sequence alignments

The amino acid sequences of trypanosome ENOs were obtained by protein BLAST using TriTrypDB [47] by using *Tbr*ENO as a query sequence.

### Crystallization, data collection and processing, and structure determination

The *Tbr*ENO - sdAbR1-10 complex (molar ratio of 2 sdAbR1-10 copies per *Tbr*ENO dimer) was purified by SEC on a ENrich SEC 650 10 x 300 (Bio-Rad) in buffer A as described above. The complex was concentrated to 8 mg ml^-1^ using a 5,000 molecular weight cut-off concentrator (Sartorius Vivaspin20) and dialyzed into buffer B (50 mM Tris-HCl, 5 mM MgCl_2_, pH 8.0) prior to setting up the crystal plates. Crystallization conditions were screened by hand or using the Mosquito Xtal 3 robot (SPT Labtech). Manual screening was performed using the hanging-drop vapor-diffusion method in 48-well plates (Hampton VDX greased) with drops consisting of 2 *µ*l protein solution and 2 *µ*l reservoir solution equilibrated against 100 *µ*l reservoir solution, while automated screening was performed via sitting drop in 96 wells plates with drops consisting of 100 nl protein solution and 100 nl reservoir solution equilibrated against 100 *µ*l reservoir solution. Commercial screens from Hampton Research (Crystal Screen, Crystal Screen 2, Crystal Screen Lite, Index, Crystal Screen Cryo), Molecular Dimensions (MIDAS, JCGS+), and Jena Bioscience (JBScreen Classic 1-10) were used for initial screening. The affinity tags of both TbrENO and sdAbR1-10 were retained for crystallization. The crystal plates were incubated at 20^°^C. Diffraction-quality crystals of *Tbr*ENO - sdAbR1-10 were obtained in JBScreen Classic 3 (Jena Bioscience) condition no. A3 (10% (w/v) PEG 4000, 10% (w/v) 2-propanol, 100 mM tri-sodium citrate, pH 5.6) and the crystals grew after approximately 7 days.

The *Tbr*ENO - sdAbR1-10 crystals were cryocooled in liquid nitrogen with the addition of 25% (v/v) glycerol to the mother liquor as a cryoprotectant in 5% increments. Data sets were collected at the ESRF synchrotron (Grenoble, France) on the ID30A-3 beamline. The best data set were processed with XDSME [48, 49]. The quality of the collected data sets was verified by close inspection of the XDS output files and through *phenix*.*xtriage* in the PHENIX package [50]. Twinning tests were also performed by *phenix*.*xtriage*. Analysis of the unit cell contents was performed with the program MATTHEWS COEF, which is part of the CCP4 package [51]. The structure of the *Tbr*ENO - sdAbR1-10 complex was determined by molecular replacement with PHASER-MR [52]. The following search models were employed for molecular replacement: i) two copies of the structure of the *Tbr*ENO dimer (PDB ID: 1OEP, [10]), and ii) four copies of an AlphaFold [53, 54] model of sdAbR1-10 (of which the CDR1 and CDR3 were removed prior to MR due to poor pLDDT scores). This provided a single solution (top TFZ = 36 and top LLG = 11365). Refinement cycles using the maximum likelihood target function cycles of *phenix*.*refine* [50] were alternated with manual building using Coot [55]. The final resolution cut-off was determined through the paired refinement strategy [56], which was performed on the PDB REDO server [57]. The crystallographic data for the *Tbr*ENO - sdAbR1-10 structure are summarized in Table 1 and have been deposited in the PDB (PDB ID: 9SLQ). Molecular graphics and analyses were performed with UCSF ChimeraX [58].

**Table 1.** Data collection and refinement statistics. Statistics for the highest resolution shell are shown in parentheses.

|  | *Tbr*ENO - sdAbR1-10 |
| --- | --- |
| **Data collection statistics** |  |
| Wavelength (Å) | 0.9677 |
| Resolution range (Å) | 47.09 - 2.33 (2.35 - 2.33) |
| Space group | 4 (P<sub>2</sub><sub>1</sub>) |
| a,b,c (Å) | 99.64 79.02 161.33 |
| α, β, γ (°) | 90 106.68 90 |
| Mosaicity (°) | 0.075 |
| Total number of measured reflections | 360734 (58859) |
| Unique reflections | 101302 (16191) |
| Multiplicity | 3.56 (3.63) |
| Completeness (%) | 98.1 (97.8) |
| < I/σ(I) > | 6.42 (0.69) |
| Wilson B-factor (Å<sup>2</sup>) | 40.44 |
| R<sub>meas</sub> (%) | 15.7 (154.5) |
| CC<sub>1/2</sub> (%) | 99.3 (67.2) |
| A.U. contains | 4 *Tbr*ENO - sdAbR1-10 complexes |
| **Refinement statistics** |  |
| CC* | 0.997 (0.586) |
| R<sub>work</sub> (%) | 0.1957 (0.4797) |
| R<sub>free</sub> (%) | 0.2428 (0.5040) |
| Number of non-hydrogen atoms | 17517 |
| macromolecules | 16519 |
| ligands | 155 |
| solvent | 843 |
| RMS bond lengths (Å) | 0.008 |
| RMS bond angles (°) | 1.05 |
| Ramachandran plot |  |
| favored (%) | 98.99 |
| allowed (%) | 0.83 |
| outliers (%) | 0.18 |
| Average B-factor (Å<sup>2</sup>) | 45.08 |
| PDB ID | 9SLQ |

### Enzyme-linked immunosorbent assay

A microtiter plate was coated overnight at 4^°^C with 1 *µ*g ml^-1^ of *Tbr*ENO, *Tco*ENO, *Tvi* ENO, or human ENO. Afterward, any unbound protein was removed by washing the plates five times with PBS containing 0.01% Tween-20 (PBST). To block residual protein-binding sites, 300 *µ*l of blocking buffer (5% milk powder in PBS) was added to each well and incubated for 2 h at room temperature (RT). Following blocking, the plate was washed three times with PBST. Then, 100 *µ*l of sdAbR1-10, prepared at a concentration of 1 *µ*g ml^-1^, was added to the blocked wells and incubated for 1 h at RT. Afterward, the plate was washed five times with PBST, and 100 *µ*l of mouse anti-HA antibody (Thermo Scientific), diluted 1:4000 in PBS, was added to each well and incubated for 1 hour at RT. The plate was then washed five times with PBST and incubated for 1 hour at RT with 100 *µ*l of streptavidin-HRP (BioLegend) diluted 1:4000 in PBS. After five final washes with PBST, 100 *µ*l TMB substrate was added to each well and incubated for 10 minutes at RT. The reaction was stopped by adding 50 *µ*l 1 M H_2_SO_4_. Finally, the absorbance as measured at a wavelength of 450 nm using a Varioskan LUX multimode microplate reader (Thermo Scientific). The data were analysed with an ordinary one-way ANOVA (GraphPad Prism).

## Results and Discussion

### sdAbR1-10 inhibits the enzymatic activity of *Tbr*ENO

We previously identified *Tbr*ENO as a potential biomarker for the detection of *T. brucei evansi* infections using camelid sdAbs [8]. Because the target of these sdAbs is a trypanosomal glycolytic enzyme, we sought to investigate whether they have the potential to inhibit *Tbr*ENO enzymatic activity. Interestingly, sdAbR1-10 appears to be a potent enzyme inhibitor as increasing concentrations severely reduce (and even completely abolish) enzymatic activity (Figure 2A).

**Fig 2.**
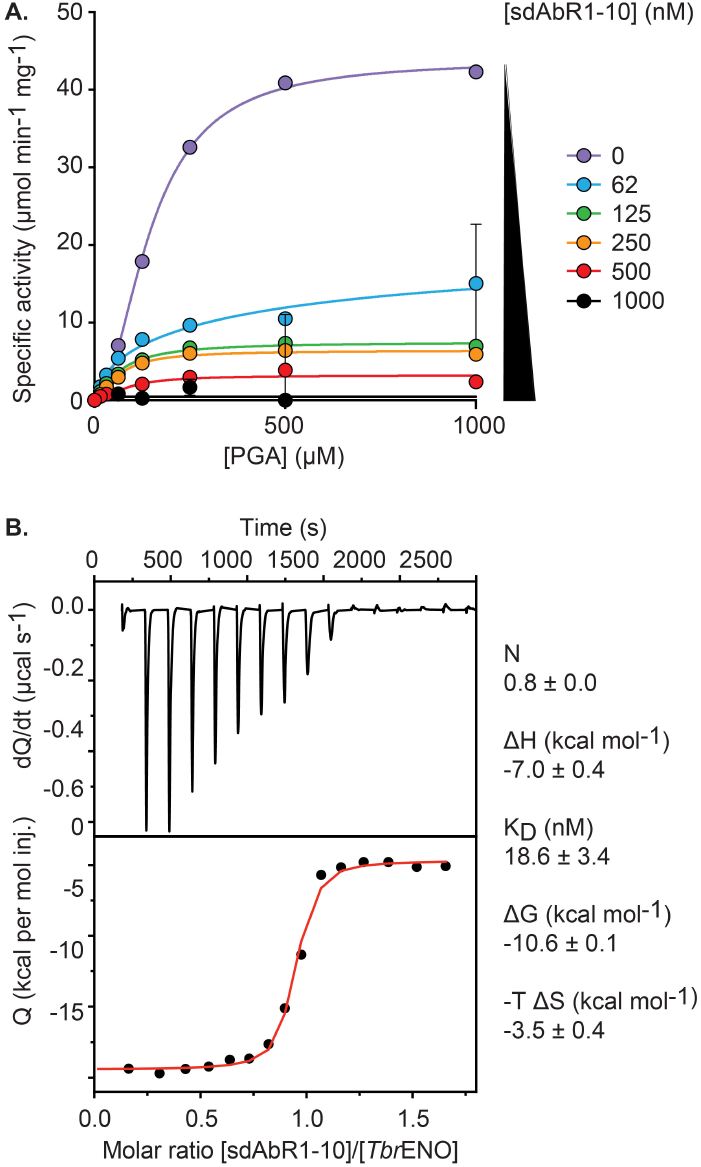
Impact of sdAbR1-10 binding on *Tbr*ENO enzyme activity and determination of the binding affinity. (A.) Enzymatic activity of recombinantly produced *Tbr*ENO at different added concentrations of sdAbR1-10 (indicated by a color code). The colored dots and traces display the experimental data points and fit, respectively. (B.) ITC measurements at 25^°^C for the binding of sdAbR1-10 to *Tbr*ENO. The top panel represents the thermogram in which the black line depicts the raw data. The bottom panel shows the isotherm. The black dots display the experimental data points, and the red trace shows the fit. The thermodynamic parameters determined via ITC data analysis are also shown.

To better understand this observation, we determined both the affinity and stoichiometry of the *Tbr*ENO - sdAbR1-10 interaction. This was first probed by isothermal titration calorimetry (ITC, Figure 2B). The resulting thermogram reveals that complex formation is exothermic and mainly enthalpically driven, of which the latter is typically the case for antibody-antigen interactions [59–62]. Analysis of the binding isotherm shows that sdAbR-10 binds its target with high nanomolar affinity (K_*D*_∼ 20 nM). Furthermore, the binding stoichiometry is close to 1, which suggests that the *Tbr*ENO homodimer contains two binding sites for sdAbR1-10 considering that the *Tbr*ENO concentration is expressed in monomer equivalents in this experiment.

The 2:2 stoichiometry of the *Tbr*ENO - sdAbR1-10 complex was further validated via a titration experiment monitored through analytical SEC. *Tbr*ENO and sdAbR1-10 were mixed in *Tbr*ENO:sdAbR1-10 molar ratios expressed in *Tbr*ENO monomer equivalents (2:1, 2:2, 2:3, and 2:4). The elution profiles of the resulting complexes were compared to the chromatograms collected for *Tbr*ENO and sdAbR1-10 alone, which expectedly elute as a dimer and monomer, respectively (Figure 3). The addition of sdAbR1-10 at 2:1 and 2:2 molar ratios cause the *Tbr*ENO elution peak to shift to the left and increase in intensity. These observations i) support the formation of a high-affinity complex (which would otherwise dissociate on the SEC column) and ii) indicate the formation of a larger *Tbr*ENO - sdAbR1-10 complex, which is further evidenced by SDS-PAGE analysis of the fractions under the elution peak. The intensity and position of the latter remain constant at 2:3 and 2:4 molar ratios, thereby showing that the *Tbr*ENO - sdAbR1-10 complex no longer increases in size at molar ratios exceeding a 2:2 stoichiometry.

**Fig 3.**
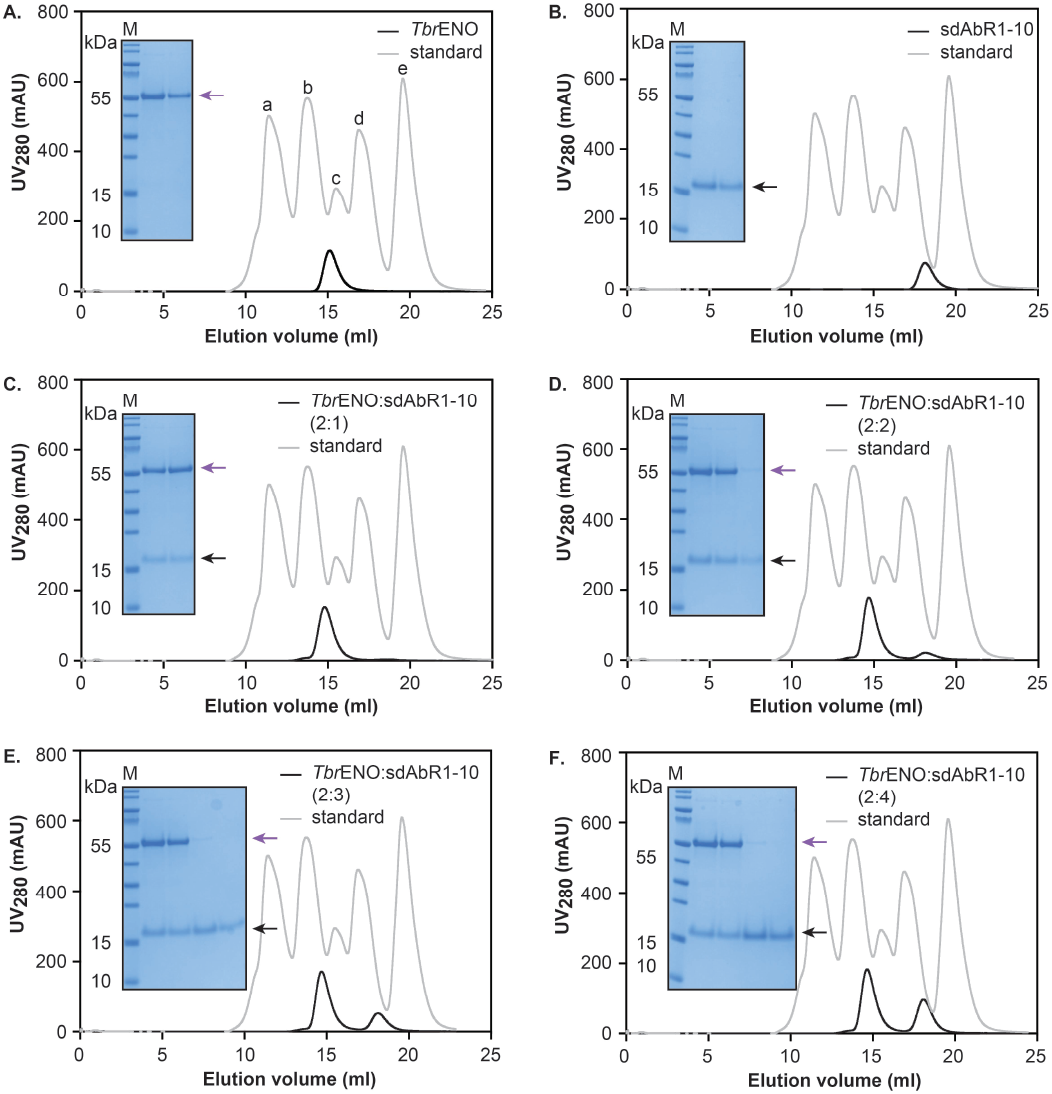
Investigation of *Tbr*ENO - sdAbR1-10 complex formation via analytical SEC. Results are shown for *Tbr*ENO alone (A.), sdAbR1-10 alone (B.), and TbrENO - sdAbR1-10 complexes prepared at various stoichiometric ratios (C. to F.. All experiments were performed on an ENRICH 650 10/30 column. The black and grey traces represent the chromatograms of the different protein samples and the BioRAD gel filtration standard, respectively. In all figures, the inset shows an SDS-PAGE analysis of the elution peaks. *Tbr*ENO (MM = 48.4 kDa) and sdAbR1-10 (MM = 14.8 kDa) are indicated by the purple and black arrows, respectively. Lane M, PageRuler Prestained Protein Ladder.

Instead, these chromatograms display an elution peak at the position expected for sdAbR1-10 alone, which represents a sdAbR1-10 excess. Hence, the analytical SEC data indeed confirm that *Tbr*ENO contains two binding sites for sdAbR1-10.

Together, these data indeed explain the inhibition data (Figure 2A). Under the employed experimental conditions and considering a high-affinity binding with a 1:1 stoichiometry, sdAbR1-10 concentrations of 62 nM or higher would lead to bound *Tbr*ENO fractions of at least 80%. The observation that these sdAbR1-10 concentrations reduce enzyme activity by percentages in the same order of magnitude suggests that each *Tbr*ENO - sdAbR1-10 binding event leads to enzyme inhibition, making sdAbR1-10 a highly potent *Tbr*ENO inhibitor.

### sdAbR1-10 inhibits *Tbr*ENO enzymatic activity by preventing closure of ‘loop 2’ and ‘loop 3’

To understand the molecular basis for the inhibition of *Tbr*ENO by sdAbR1-10, the high-resolution structure of the *Tbr*ENO - sdAbR1-10 complex was determined via MX. A general overview of the crystal structure reveals that two sdAbR1-10 molecules are bound to the *Tbr*ENO dimer (Figure 4A), which is in accordance with the in-solution stoichiometry determined via analytical SEC and ITC. The epitope is located on *Tbr*ENO’s C-terminal eightfold *α*/*β* barrel domain and displays no direct overlap with the active site. A detailed interaction analysis reveals that the sdAb contacts *Tbr*ENO ‘loop 2’ and ‘loop 3’ residues through both its complementarity determining regions and framework regions (CDRs and FRs, respectively; Table 2 and Figure 4, panels B and C). Water plays an especially important role in the sdAb-antigen recognition event as advocated by the multitude of water-mediated hydrogen bonds observed at the interaction interface, which is again consistent with the existing literature on the molecular aspects of antibody-antigen interactions [59–61, 63].

**Table 2.**
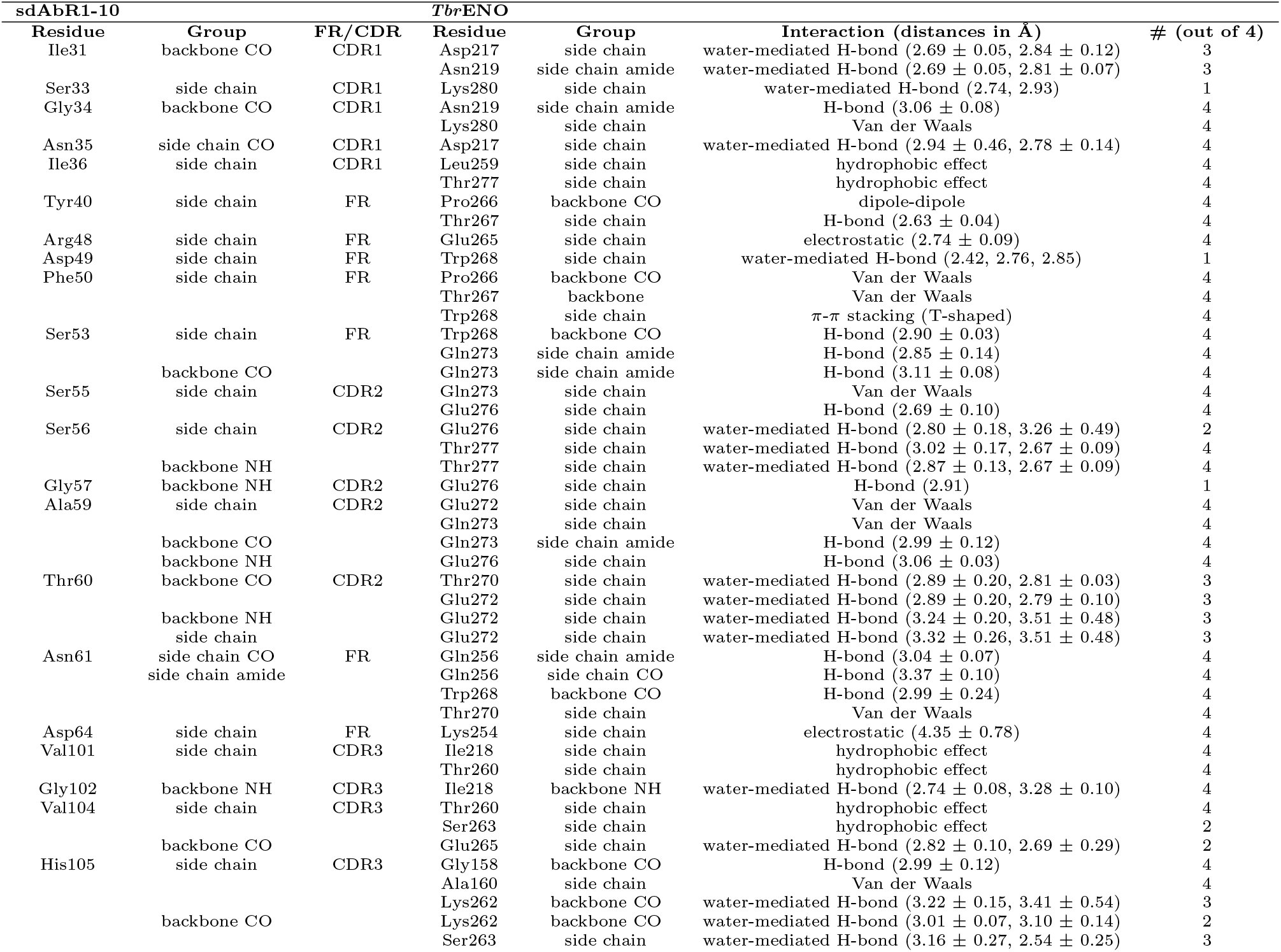
List of interactions between sdAbR1-10 and *Tbr*ENO. The # symbol indicates the number of times the interaction was observed over the total of four *Tbr*ENO - sdAbR1-10 complexes present in the asymmetric unit. The average distances are only given for hydrogen bonds or electrostatic interactions.

**Fig 4.**
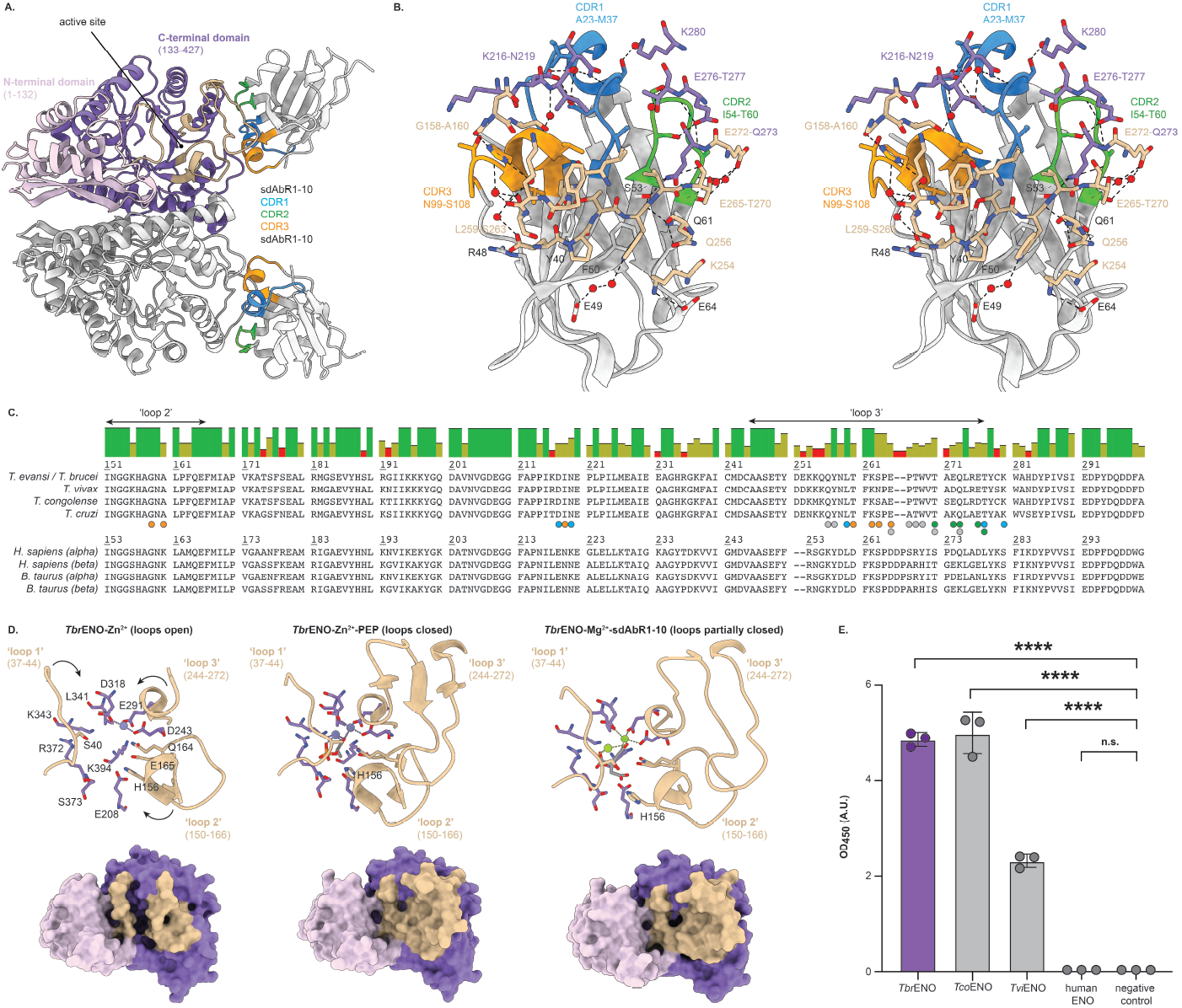
Structural basis for the selectivity of sdAbR1-10 towards trypanosomal ENOs. (A.) Cartoon representation of the *Tbr*ENO - sdAbR1-10 complex observed in the crystal, in which one *Tbr*ENO dimer is bound by two copies of sdAbR1-10. The *Tbr*ENO dimer is colored as in Figure 1A. sdAbR1-10 is depicted in light grey and its CDRs are colored in blue (CDR1), green (CDR2), and orange (CDR3). (B.) Stereo view of the interactions at the *Tbr*ENO - sdAbR1-10 interface. sdAbR1-10 is depicted in cartoon representation as in (A.). For reasons of clarity, only the *Tbr*ENO residues that are part of the epitope are shown and colored as in (A). All interacting residues are labelled and shown in stick representation. Hydrogen bonds and salt bridges are indicated by black dashed lines. (C.) Sequence alignment between trypanosomal, human, and bovine ENOs. The colored bars above the sequence alignment represent the percentage of sequence identity: green (100%), green-brown(between 30% and 100%), and red (below 30%). The residues marked by the colored circles are contacted by sdAbR1-10 according to the color code in (A.): CDR1 (blue), CDR2 (green), CDR3 (orange), and FR (light grey). (D.) Comparison of *Tbr*ENO structures with different active site loop conformations as presented in in Figure 1B: *Tbr*ENO-Zn^2+^ (loops open, PDB ID 2PTW [11]), *Tbr*ENO-Zn^2+^-PEP (loops closed, 2PTY [11]), and *Tbr*ENO-Mg^2+^-sdAbR1-10 (loops partially closed, PDB ID 9SLQ, this work). (E.) ELISA results demonstrating that sdAbR1-10 only binds trypanosomal ENOs, and not to human ENO. The mean values with standard deviation of three independent measurements are shown. Statistical significance was determined by one-way analysis of variance; \*\*\*\**p ≤* 0.0001, n.s. = not significant.

Comparing the conformations of the three ‘loops’ surrounding the enzyme’s active site in the sAbR1-10 - bound complex to *Tbr*ENO structures in which all three ‘loops’ are completely opened or closed reveals an interesting disparity: while ‘loop 1’ adopts a closed structure, both ‘loop 2’ and ‘loop 3’ reside in an open conformation (Figure 4D). The closing of ‘loop 1’ can be explained by the binding of a citrate molecule to the active site due to its presence in the crystallisation cocktail. Here, citrate acts as a substrate mimic as two of its carboxylic acid groups engage in interactions that would normally be provided by the carboxylic acid and phosphoryl groups of D-2-phosphoglycerate/PEP. In contrast, ‘loop 2’ and ‘loop 3’ remain open, which appears to be the consequence of sdAbR1-10 contacting ‘loop 2’ and mainly ‘loop 3’ residues. Interestingly, while no electron density could be observed for ‘loop 3’ in the *Tbr*ENO crystal structure containing open ‘loops’, clear electron density could be discerned for this region in the *Tbr*ENO - sdAbR1-10 complex, thereby indicating ‘loop 3’ stabilisation in the latter. Hence, sdAbR1-10 binding would appear to stabilise both ‘loop 2’ and ‘loop 3’ in an open conformation. In turn, this would prevent them from completely closing, which would indeed impede successful active site formation and thus lead to an inhibition in catalytic activity. Interestingly, to the best of our knowledge, this is the first report of a potent, specific *Tbr*ENO inhibitor that abolishes enzyme activity by binding to a site other than the enzyme’s active site.

### The sdAbR1-10 epitope is conserved in trypanosomal, but not human and bovine enolases

A sequence comparison of trypanosomal, and human and bovine ENOs reveals that the epitope targeted by sdAbR1-10 is conserved within trypanosomatids, but not in human and bovine ENOs (Figure 4C). Hence, sdAbR1-10 would only be expected to bind trypanosomal ENOs, but not their human and bovine counterparts. To confirm this hypothesis experimentally, the binding of sdAbR1-10 to various ENOs was tested through an enzyme-linked immunosorbent assay (ELISA; Figure 4E). Indeed, while a clear binding signal could be detected for interaction with ENOs from *T. brucei, T. congolense*, and *T. vivax*, no binding signal could be observed against human ENO. We could not obtain bovine ENO for experimentation, but given a sequence identity of at least 85 % between human and bovine ENOs, we believe the observed outcome will be the same. Based on these results and our experience with an sdAb targeting trypanosomal pyruvate kinase [62], we assume that sdAbR1-10 will also inhibit the activities of *T. congolense* and *T. vivax* ENOs, although this has not been experimentally confirmed. We anticipate that the inhibition of other trypanosomal ENOs by sdAR1-10 will occur with reduced efficiency given that the targeted epitope is not 100% conserved. In our previous work with an sdAb targeting trypanosomal pyruvate kinase [62], we observed that subtle changes in the amino acids belonging to the targeted epitope lead to a reduction in binding affinity, and that this correlates with a decrease in inhibition efficiency. Finally, given that sdAbR1-10 exhibits no binding to human ENO, we expect that this sdAb will have no impact on its enzymatic activity.

## Conclusion

In this paper, we have reported the discovery of a camelid sdAb (sdAbR1-10) that specifically recognises trypanosomal (but not host) ENOs. sdAbR1-10 unexpectedly turned out to be a specific and potent *Tbr*ENO inhibitor, and its mechanism of action was thoroughly studied through a combination of protein biochemistry, biophysics, and structural biology. Interestingly, this is the second time that we serendipitously discover an sdAb (initially generated within a diagnostic context) that displays potent inhibitory properties against its target enzyme antigen [62]. In both cases, enzyme activity is abolished through a mechanism that does not involve direct binding of the enzyme’s active site. While it remains unclear to us at this stage whether such mechanisms could potentially be exploited for the design of novel chemotherapeutics, reporting these findings may be of particular interest to those active in this line of research.

## Acknowledgments

This work was supported by a research grants of the University of Antwerp (DOCPRO1 - FFB190197 and ‘Bijzonder Onderzoeksfonds’ 41391 awarded to Y.G.-J.S.), and the ‘Fonds voor Wetenschappelijk Onderzoek - Vlaanderen’ (FWO-Vlaanderen, G013518N to S.M.). The authors wish to thank the staff of the ESRF synchrotron (Philippe CARPENTIER) for outstanding beam line support. The authors declare to have no conflict of interest.

